# Protein-independent regulation by transgene-derived small interfering RNAs rewires endogenous regulatory networks to enhance plant growth, architecture, and drought performance

**DOI:** 10.64898/2026.09.08.750206

**Authors:** Gustavo J. Vannay, Joaquín E. García, José P. Murguía, María Lourdes Bruno, Luciano Caraballo, Elina Welchen, Damián A. Cambiagno, Raquel L. Chan, Matías Capella

## Abstract

Plants produce diverse small interfering RNA (siRNA) molecules that modulate development, environmental responses, and immunity. Although transgene-derived siRNAs are traditionally viewed as mediators of gene silencing, whether they can actively regulate endogenous host pathways remains largely unexplored. Previously obtained Arabidopsis, wheat, and soybean plants expressing the sunflower gene encoding the transcription factor HaHB4 exhibited water deficit tolerance. Here, we show that expressing inverted-repeat constructs that generate *HaHB4*-derived siRNAs without producing the HaHB4 protein bypasses transgenic growth penalties and instead enhances vegetative vigor and reproductive performance. In Arabidopsis, these DCL-dependent siRNA-producing lines exhibited enhanced root growth, increased stem and pith areas, increased cauline branching, and higher seed yield under both optimal and water-limiting conditions. Transcriptomic analysis revealed convergent repression of biotic stress-related genes, accompanied by increased bacterial susceptibility and reduced sensitivity to salicylic acid-mediated growth inhibition, suggesting an altered balance between immunity and growth. Functional characterization of candidate endogenous HD-Zip I targets further showed that *athb20* and *athb53* mutants recapitulated the increased stem expansion and cauline branching of the RNAi lines, respectively, pointing to endogenous HD-Zip I genes as candidate mediators of these traits. Remarkably, these effects were observed in newly obtained transgenic soybean plants, where expression of *HaHB4*-derived siRNAs enhanced vegetative vigor under controlled growth conditions. Overall, these findings show that transgene-derived siRNAs act independently of protein function to rewire endogenous regulatory networks, providing a potential strategy to optimize crop architecture and yield.

## Introduction

Ensuring global food security remains a major challenge as rapid population growth concurs with climate warming, increased environmental instability, and water scarcity. To mitigate these threats, significant advances have been made through the use of biotechnological tools, which have enabled sustainable increases in both seed yield and abiotic stress tolerance in major crops, including wheat, maize, rice, and soybean (Riaz et al., 2025). Notably, most of these biotechnological strategies leverage endogenous molecular pathways that plants possess to cope with adverse environmental conditions (Villalobos-López et al., 2022).

Key players in these adaptive responses are transcription factors (TFs), which function as master regulators of gene expression to orchestrate developmental plasticity and facilitate adaptation to environmental challenges. Given their capacity to modulate complex transcriptomic networks, TFs are powerful tools for enhancing agronomic traits such as seed yield, stress tolerance, and nutrient acquisition (Bhoite et al., 2025; Wang et al., 2016). A prominent example of a TF-based strategy to stabilize crop production under water deficit is the HB4® technology, the market name for a genetic construct carrying the sunflower *HaHB4* (*<u>H</u>elianthus <u>a</u>nnuus <u>H</u>omeo<u>b</u>ox <u>4</u>*), which encodes a transcription factor in the homeodomain-leucine zipper (HD-Zip) family (González et al., 2019; Ribichich et al., 2020). Plants bearing this technology are already approved in several countries (www.isaaa.org) and commercialized. Members of this plant-exclusive family are defined by the presence of a DNA-binding domain named the homeodomain (HD), coupled with a leucine zipper (Zip) that mediates homo- or heterodimerization. HD-Zip proteins are categorized into four distinct subfamilies based on sequence homology within the HD-Zip domain, gene structure (intron/exon organization), the presence of additional conserved motifs, and their specific regulatory roles (Capella et al., 2016). Although the HD-Zip I subfamily has traditionally been recognized as a pivotal contributor to abiotic stress responses, recent insights underscore its involvement in diverse developmental programs and hormonal signaling networks (Perotti et al., 2021, 2017).

Extensive research has established that transgenic Arabidopsis plants expressing the sunflower *HaHB4* gene exhibit enhanced water deficit tolerance by modulating ethylene sensitivity and delaying senescence (Dezar et al., 2005; Manavella et al., 2006). HaHB4 also governs additional signal transduction pathways, such as the induction of jasmonic acid, which in turn fortifies defenses against herbivory, and the repression of genes associated with photosynthesis during periods of darkness (Manavella et al., 2008a, 2008b). *HaHB4*- expressing soybean and wheat consistently outperform their wild-type counterparts, due to an increased grain number. Particularly under fluctuating water availability and high temperatures during critical reproductive phases, *HaHB4*-transgenic plants exhibited less yield penalty than their controls (Ayala et al., 2026; González et al., 2019; Ribichich et al., 2020). In wheat, this important trait is particularly expressed when the thermal challenge occurs during the pre-anthesis stage (Murguía et al., 2026a).

Like viral DNA and transposons, transgene overexpression often generates aberrant RNA molecules that might trigger a repression mechanism known as posttranscriptional gene silencing (PTGS) (Béclin et al., 2002; Dalmay et al., 2000). Aberrant transcripts are recognized by the RNA-dependent RNA polymerase RDR6, which generates double-stranded RNAs (dsRNA), which are subsequently processed into 20-22 nucleotide RNA duplexes by Dicer family proteins, such as DCL2 (Dicer-like 2) and DCL4 (Henderson et al., 2006; Luo and Chen, 2007; Mourrain et al., 2000; Parent et al., 2015). One strand of the small RNAs is finally loaded into the Argonaute 1 (AGO1)-containing RNA-induced silencing complex (RISC), resulting in mRNA cleavage or translational inhibition (Marí-Ordóñez et al., 2013; Rössner et al., 2022). If loaded into AGO6 rather than AGO1, the siRNA-protein complex is targeted to specific genomic loci via RNA Polymerase V (Pol V) scaffolding transcripts (McCue et al., 2015). This interaction recruits the methyltransferase DOMAINS REARRANGED METHYLTRANSFERASE 2 (DRM2) to catalyze de novo DNA methylation and silence Pol II transcription. Once established, this epigenetic modification is maintained by the canonical RNA-dependent DNA methylation (RdDM) pathway, where RDR2 generates dsRNA from Pol IV templates, which is then processed by DCL3 and loaded into AGO4 (or AGO2) to recruit DRM2 (Cuerda-Gil and Slotkin, 2016). Additionally, RDR6-derived dsRNA can directly feed into the RdDM pathway if cleaved by DCL3 into 24-nt siRNAs (Marí-Ordóñez et al., 2013).

HD-Zip I encoding genes are no exception, and their overexpression can be silenced by the generation of siRNA. For instance, the heterologous overexpression of coffee *HB12* results in detectable transgene-derived siRNA (Cruz et al., 2024). Overexpression of endogenous HD-Zip I genes in an *rdr6* background effectively prevented gene silencing, resulting in plants with serrated leaves and enhanced tolerance to water deficit, a phenomenon likely attributable to increased protein levels (Miguel et al., 2020; Ribone et al., 2017; Romani et al., 2016). Therefore, careful consideration is warranted when overexpressing transgenes to improve plants, as PTGS mechanisms may hinder the intended effects and potentially lead to unintended consequences. Nevertheless, whether the consequent generation of transgene-derived small RNAs might influence global gene expression and ultimately confer beneficial traits remains largely unexplored.

Here, we demonstrate that the expression of *HaHB4*-derived siRNAs enhances vegetative and reproductive performance in Arabidopsis. Plants accumulating such siRNAs exhibited increased leaf area, thicker stems, and more number of cauline branches than controls, resulting in an increase in seed yield. Furthermore, these lines exhibited better performance under water-deficit conditions. Transcriptomic profiling revealed the downregulation of numerous biotic stress-related genes, rendering the plants more susceptible to *Pseudomonas syringae* infection and attenuating sensitivity to salicylic acid-mediated growth inhibition. Furthermore, the accumulation of *HaHB4*-derived siRNAs might negatively influence the expression of several endogenous HD-Zip I genes, likely driving some of the observed morphological and physiological changes. Transgenic soybean lines expressing *HaHB4*-derived siRNAs exhibited increased hypocotyl and epicotyl width, expanded leaf area, and higher shoot biomass, demonstrating cross-species conservation of these beneficial traits. Collectively, our findings demonstrate that transgene-derived small RNAs can serve as powerful tools modulating global transcriptional landscapes and improving agronomic traits.

## Results

### High expression levels of the *HaHB4* transgene negatively affect plant development

Given that the expression of endogenous genes encoding HD-Zip I transcription factors is susceptible to transgene-mediated silencing in Arabidopsis (Miguel et al., 2020; Ribone et al., 2017; Romani et al., 2016), we investigated whether a similar post-transcriptional mechanism acts upon ectopic *HaHB4* expression. Elevated *HaHB4* transcript levels in Arabidopsis typically resulted in rounded leaves, shortened petioles, compact rosettes, and delayed flowering (Cabello et al., 2007; Dezar et al., 2005). To evaluate the frequency of such high-expression phenotypes, we introduced the genetic construct *35S:HaHB4* (Dezar et al., 2005) into Col-0 wild-type (WT) plants. Notably, we observed that approximately 20% of the resulting independent transformants exhibited the characteristic small-plant phenotype (Figure 1a,b).

**Figure 1.**
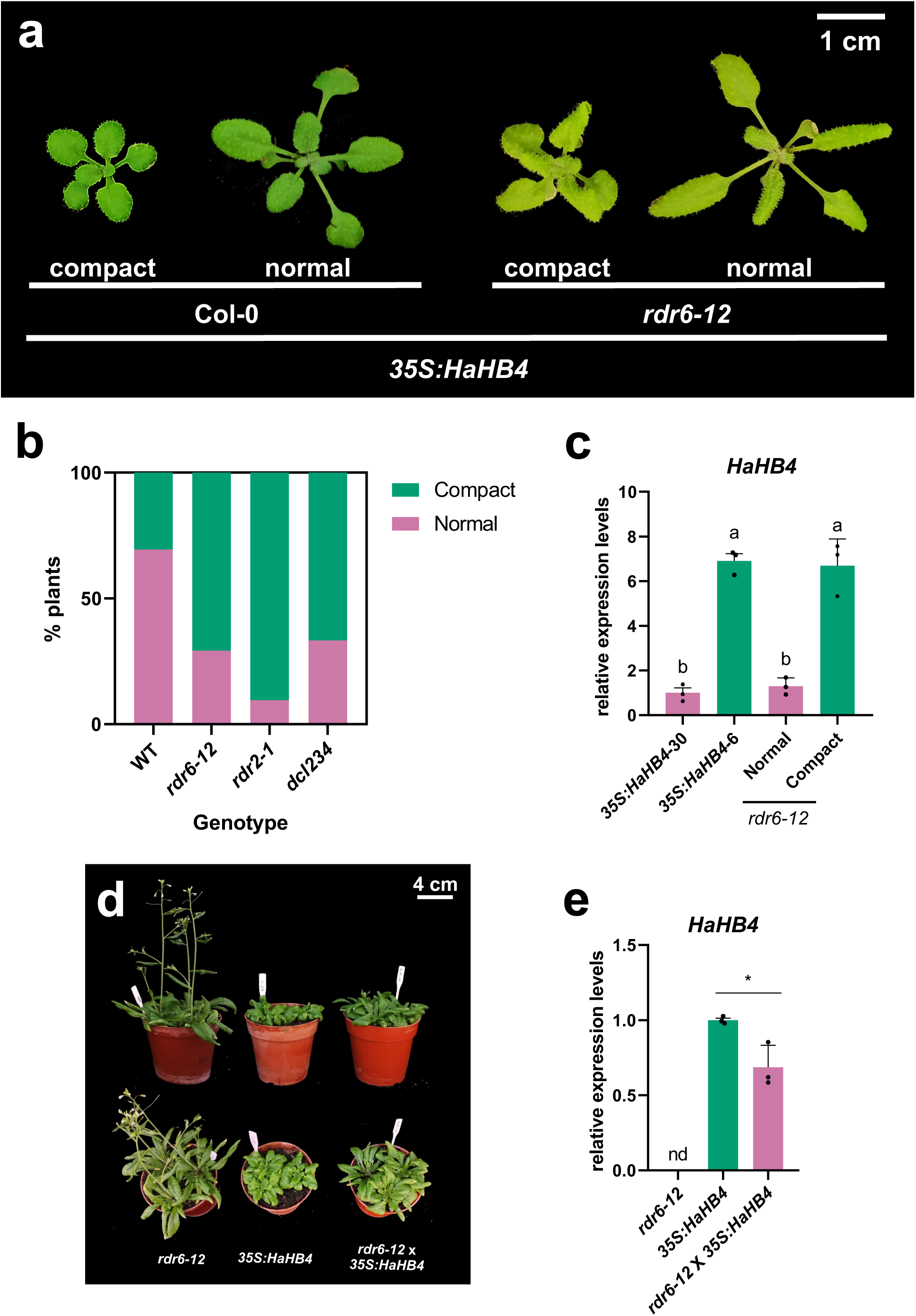
***HaHB4* expression is silenced by the small RNA machinery in Arabidopsis.** (a) Representative images of Col-0 and *rdr6-12* plants transformed with the *35S:HaHB4* construct, illustrating the wild-type (normal) and small-plant (compact) phenotypes. Scale bar: 1 cm. (b) Proportion of 20-day-old Col-0, *rdr6-12*, *rdr2-1*, and *dcl234* plants transformed with the *35S:HaHB4* construct displaying normal or compact phenotypes. Pink and green bars represent the percentages of normal and compact plants, respectively. (c) *HaHB4* transcript levels quantified by RT-qPCR in three independent *rdr6-12* transgenic plants displaying either normal (pink) or compact (green) phenotypes. The low-expressing *35S:HaHB4*-30 and high-expressing *35S:HaHB4*-6 lines served as reference controls. Expression levels were normalized to *ACTIN2/8* and are shown on a linear scale relative to the *35S:HaHB4*-30 line. (d) Representative images showing 35-day-old *rdr6-12*, *35S:HaHB4*-6, and the genetic cross (*rdr6-12 35S:HaHB4*-6). Scale bar: 4 cm. (e) Quantification of *HaHB4* expression in the lines described in (d). For (c) and (e), data are means (±SEM) of *n*=3 independent biological replicates. Different letters indicate significant differences among means as determined using one-way ANOVA followed by Tukey’s *post-hoc* test (*P*<0.05).

To determine whether siRNA biogenesis underlies the silencing of *HaHB4* expression across independent transgenic events, we next assessed the impact of the *35S:HaHB4* construct when introduced into key small RNA processing-deficient backgrounds, including *rdr2-1*, *rdr6-12*, and *dcl2-1/dcl3-1/dcl4-2t* (*dcl234*). In striking contrast to WT plants, transformation of these silencing-pathway mutants yielded the small-plant phenotype in more than 80% of the descendants (Figure 1b). Transcript quantification using reverse transcription followed by quantitative real-time PCR (RT-qPCR) showed that high *HaHB4* expression correlated with the small-plant phenotype in the *rdr6-12* background (Figure 1c). Indeed, three independent transgenic *rdr6-12* lines exhibiting the small-plant phenotype accumulated *HaHB4* transcript levels comparable to those of the high-expressing *35S:HaHB4*-6 reference line (Cabello et al., 2007) (Figure 1c). In contrast, the lines displaying a WT-like morphology exhibited the low expression levels found in *35S:HaHB4*-30 plants (Figure 1c).

RDR6 is a key component in establishing post-transcriptional silencing of highly expressed sense transgenes (Dalmay et al., 2000; Mourrain et al., 2000). Once silencing is established, its maintenance can become less dependent on RDR6, depending on the transgene and the silencing mechanism involved (Béclin et al., 2002; Himber et al., 2003). To determine whether RDR6 is required to maintain *HaHB4* transgene silencing, we crossed the *rdr6-12* mutant with the *35S:HaHB4*-6 high-overexpressor line and evaluated the resulting phenotype. Notably, *rdr6-12* plants overexpressing *HaHB4* displayed smaller leaves and delayed flowering, closely resembling the parental *35S:HaHB4*-6 plants (Figure 1d). Although expression analysis showed that *HaHB4* transcript levels were reduced by half in the background relative to the *35S:HaHB4*-6 line (Figure 1e), the preservation of the developmental defects implies that phenotype maintenance is largely independent of RDR6 once sufficient *HaHB4* expression is established. Together, our findings establish that the *HaHB4* transgene is subjected to silencing in Arabidopsis and suggest that the observed phenotypic aberrations may be a direct consequence of HaHB4 protein activity.

### Expression of only *HaHB4*-derived siRNAs significantly improved plant performance

Based on these initial observations, we postulated that small RNAs derived from the *HaHB4* sequences can confer beneficial traits to Arabidopsis plants *per se*, independently of the protein. To test this hypothesis, we designed genetic constructs under the control of the *35S* promoter containing inverted repeats of two distinct *HaHB4* segments, separated by an intervening intron to facilitate the formation of hairpin RNAs (Smith et al., 2000). Specifically, we generated constructs encompassing the N-terminus and homeodomain or the leucine zipper and C-terminal regions of the *HaHB4* CDS (named thereafter *HDi* and *CTRi*, respectively) (Figure 2a). Unlike *35S:HaHB4*-30 transgenic lines, plants transformed with the *HDi* or *CTRi* constructs exhibited a vigorous growth phenotype, characterized by significantly longer hypocotyls and larger leaves, while maintaining WT-like stem length (Figure 2b-d). Furthermore, both primary root length and total lateral root (LR) length were significantly increased in seedlings expressing *HaHB4*-derived siRNAs compared to WT and *35S:HaHB4*-30 controls (Figure 2e-g). In contrast, LR density remained unaltered among all evaluated genotypes (Figure 2h).

**Figure 2.**
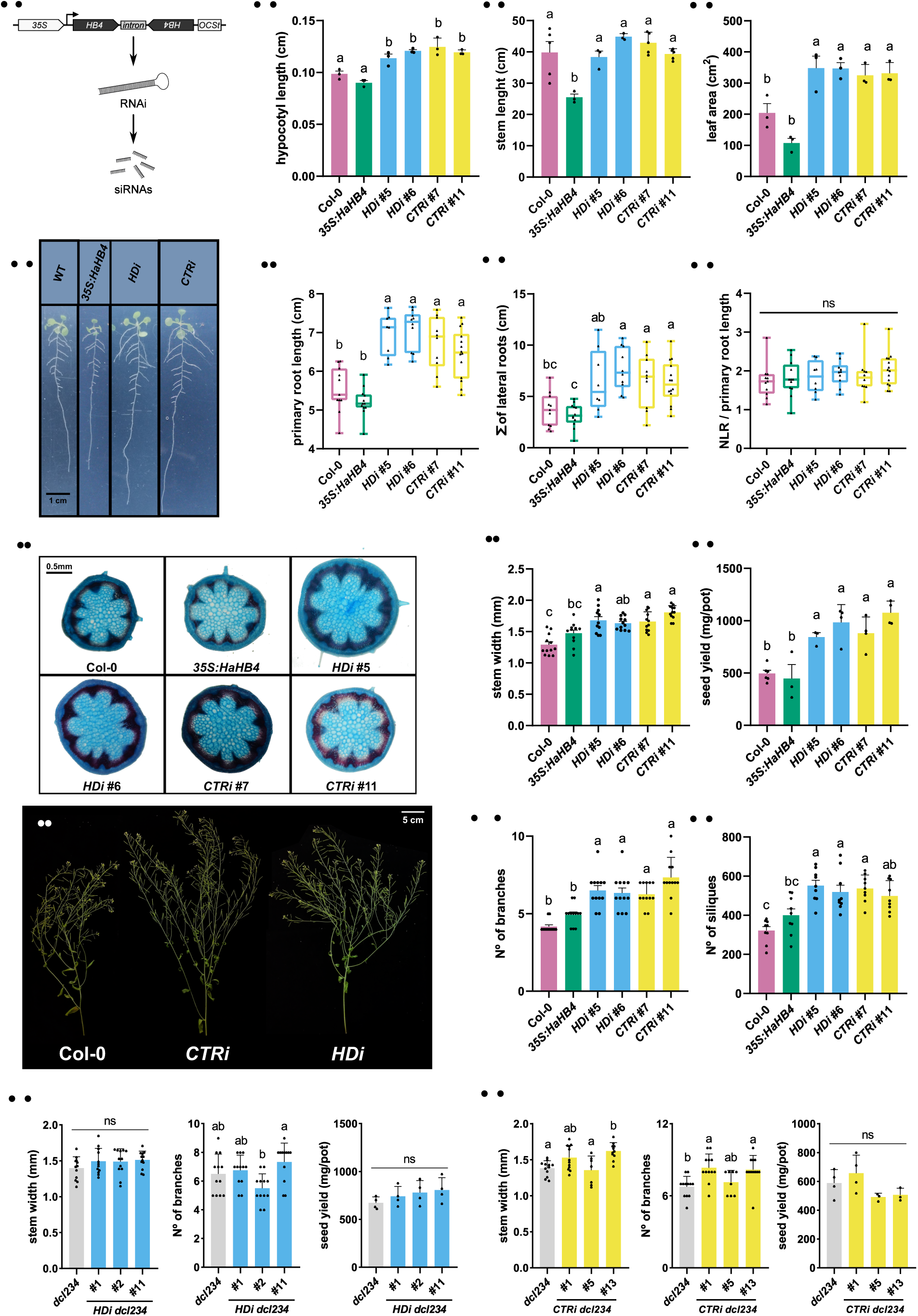
Expression of *HaHB4*-derived siRNA improves plant performance in Arabidopsis. (a) Schematic representation of the genetic constructs designed to generate small RNAs. (b) Quantification of hypocotyl length of 6-day-old Col-0, the low-expressing *HaHB4* line 30 (*35S:HaHB4*), and two independent Col-0 lines transformed with either the *HDi* or *CTRi* constructs. (c) Stem length quantification of 36-day-old Col-0, *35S:HaHB4*, and two independent *HDi* or *CTRi* lines. (d) Leaf area measurements of the plants described in (c), grown for 25 days. (e) Illustrative pictures of 10-day-old Col-0, *35S:HaHB4*, and two independent *HDi* or *CTRi* lines. Scale bar: 1 cm. (f-h) Primary root length (f), total lateral root length (g), and number of lateral roots relative to the primary root length (h) measured in the seedlings described in (e). (i) Representative transverse stem sections of the first internode from 30-day-old Col-0, *35S:HaHB4*, and two independent *HDi* or *CTRi* lines. Freehand cross-sections were stained with Astra blue-Safranin. Scale bar: 0.5 mm. (j) Stem width measured 30 days after sowing for the plants described in (c). (k) Seed yield evaluated at the end of the life cycle of the plants described in (c), expressed as mg/pot. (l) Representative pictures of the main stem from 50-day-old Col-0, and the *HDi* or *CTRi* lines. (m,n) Quantification of the number of cauline branches (m) and siliques (n) in the plants described in (j), 50 days after sowing. (o,p) Stem width, number of cauline branches, and seed yield from *dcl2-1 dcl3-1 dcl4-2t* (*dcl234*), and three independent *HDi* (o) or *CTRi* (p) lines in the *dcl234* background. For (b) to (p), seedlings or plants were grown under long-day conditions at 22 °C. For soil-filled pots, all experiments were performed with 3 or 4 pots per genotype, with three plants per pot. For (b)-(d), (f)-(g), (j), (k), and (m)-(p), data are means (±SEM) of *n*=3-12 independent biological replicates. Different letters indicate significant differences among means as determined using one-way ANOVA followed by Tukey’s *post-hoc* test (*P*<0.05). ns, not significant.

Notably, both *HDi* and *CTRi* plants also showed wider stems, a characteristic shared with *35S:HaHB4* plants (Figure 2i,j). Histological cross-section analyses revealed that these new plants exhibited larger stem and pith areas, but had the same number of vascular bundles as controls (Figure 2i; Figure S1). Consistent with the established link between stem diameter and overall yield (Cabello and Chan, 2019), we observed a substantial increase in seed production in *HDi* and *CTRi* plants compared to WT (Figure 2k). This productivity boost was a consequence of a greater number of cauline branches (CI), which concomitantly elevated total siliques (Figure 2l-n). Notably, these distinct traits of *HDi* and *CTRi* lines were also observed under short-day conditions (Figure S2). To confirm that the observed differential traits in the RNAi plants resulted from siRNA production, we introduced the RNAi constructs into a *dcl234* triple mutant background, which lacks functional dicer-like proteins and therefore cannot process double-stranded RNA into small RNAs. As expected, expression of *HDi* and *CTRi* failed to increase stem width or seed yield in the *dcl234* background (Figure 2o,p), indicating that siRNA generation is necessary for these phenotypic modifications.

### *HaHB4*-derived siRNAs enhance seed production while consuming less water under water deficit

The heterologous expression of *HaHB4* consistently improved plant performance under water-deficit conditions across various species, including Arabidopsis, soybean, and wheat (Dezar et al., 2005; González et al., 2019; Ribichich et al., 2020). Hence, we evaluated whether the expression of *HaHB4*-derived siRNAs influences seed production under water-limiting conditions. To test this, plants were cultivated under optimal watering regimes for 30 days and subsequently subjected to a 20-day water-deficit period by maintaining field capacity (FC) at 50% (Figure 3a). Following the drought treatment, the plants were fully rewatered and grown under well-watered conditions until seed harvest. Notably, despite a drought-induced yield penalty relative to water-deficit conditions, *HDi* and *CTRi* lines produced a significantly higher overall seed yield than WT controls, accompanied by an increased number of cauline branches (Figure 3b,c). In contrast, *35S:HaHB4*-30 plants displayed WT-like seed production under these water-limiting conditions (Figure 3c). It is worth noting that while *35S:HaHB4* lines are known to survive severe water-deficit stress, their reproductive yield under these specific conditions had not previously been quantified in Arabidopsis (Dezar et al., 2005). Furthermore, both *35S:HaHB4* and siRNA-expressing lines exhibited higher seed yield accompanied by lower water consumption than WT plants (Figure 3d). Consequently, the *HDi* and *CTRi* plants achieved a slightly higher harvest index compared to untransformed controls (Figure 3e). Altogether, these findings demonstrated that *HaHB4*-derived siRNAs enhanced plant growth and reproductive fitness under both normal and water-limiting conditions.

**Figure 3.**
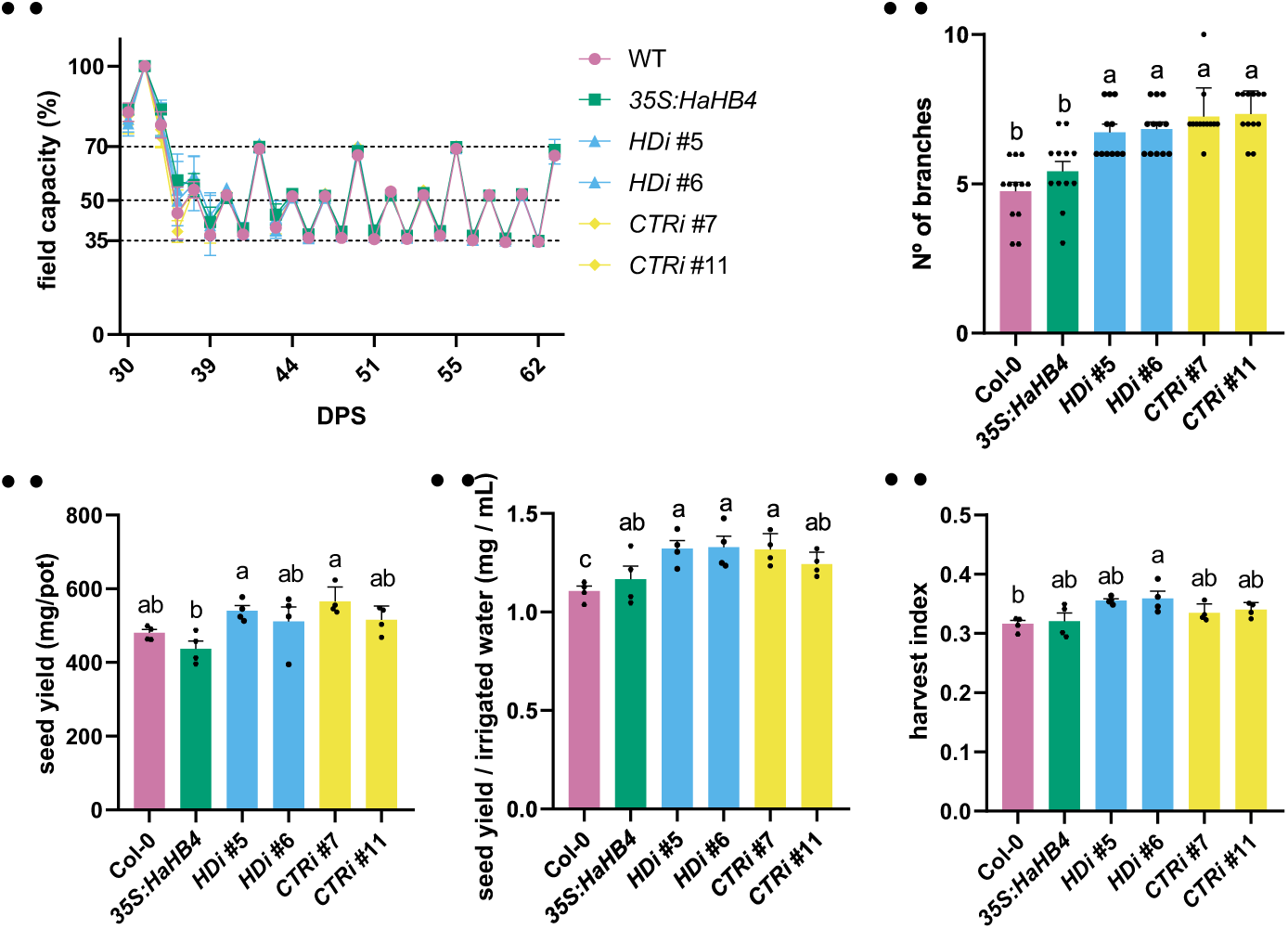
The *HDi* and *CTRi* lines outyield the WT control under water-deficit conditions. Plants were cultivated under well-watered conditions for 30 days and subsequently subjected to a 20-day water-deficit period by maintaining the field capacity at 50%. (a) Water consumption of Col-0, the low-expressing *HaHB4* line 30 (*35S:HaHB4*), and two independent *HDi* or *CTRi* lines during the water-limiting period. (c) Number of cauline branches of the plants described in (a) 50 days after sowing. (c) Seed yield evaluated at the end of the life cycle of the plants described in (a), expressed as mg/pot. (d) Seed weight related to the consumed water (SW/CW) of the plants described in (a), determined by calculating the ratio of total accumulated dry biomass to the total volume of water consumed during the water-deficit treatment. (e) Harvest index evaluated at the end of the life cycle of the plants described in (a). Data are means (±SEM) of *n*=4 independent biological replicates. Different letters indicate significant differences among means as determined using one-way ANOVA followed by Tukey’s *post-hoc* test (*P*<0.05).

### Convergent transcriptomic profiles revealed that *HDi* and *CTRi* plants impact on plant immunity

To elucidate how *HaHB4*-derived siRNAs influence plant development, we conducted a comparative RNA-Seq analysis on leaves and stems of 30-day-old WT, *HDi*, and *CTRi* plants. We identified 690 and 843 differentially expressed genes (DEGs; log_2_ fold change > 1 or < −1, *p*-adj < 0.05) in leaves and stems of *HDi* plants relative to WT controls (Figure 4a, Tables S1 and S2). Parallel analysis of *CTRi* lines detected 820 DEGs in leaves and 729 in stems compared to the WT (Figure 4a, Tables S3 and S4). Surprisingly, the majority of DEGs detected in siRNA-expressing leaves were distinct from the transcriptional changes observed in *35S:HaHB4* plants (Manavella et al., 2006) (Figure S3). Given the striking phenotypic similarities between *HDi* and *CTRi* plants, we next investigated whether these independent lines shared common transcriptional profiles. By comparing their transcriptomes, we found a significant overlap among the modulated gene sets in both leaves and stems (Figure 4b). Consistent with this observation, pairwise scatterplot comparisons showed a strong correlation between the expression changes in *HDi* and *CTRi* plants across both tissues (Figure 4c). These findings suggested that both RNAi lines trigger equivalent regulatory pathways, regardless of their different siRNA sequences.

**Figure 4.**
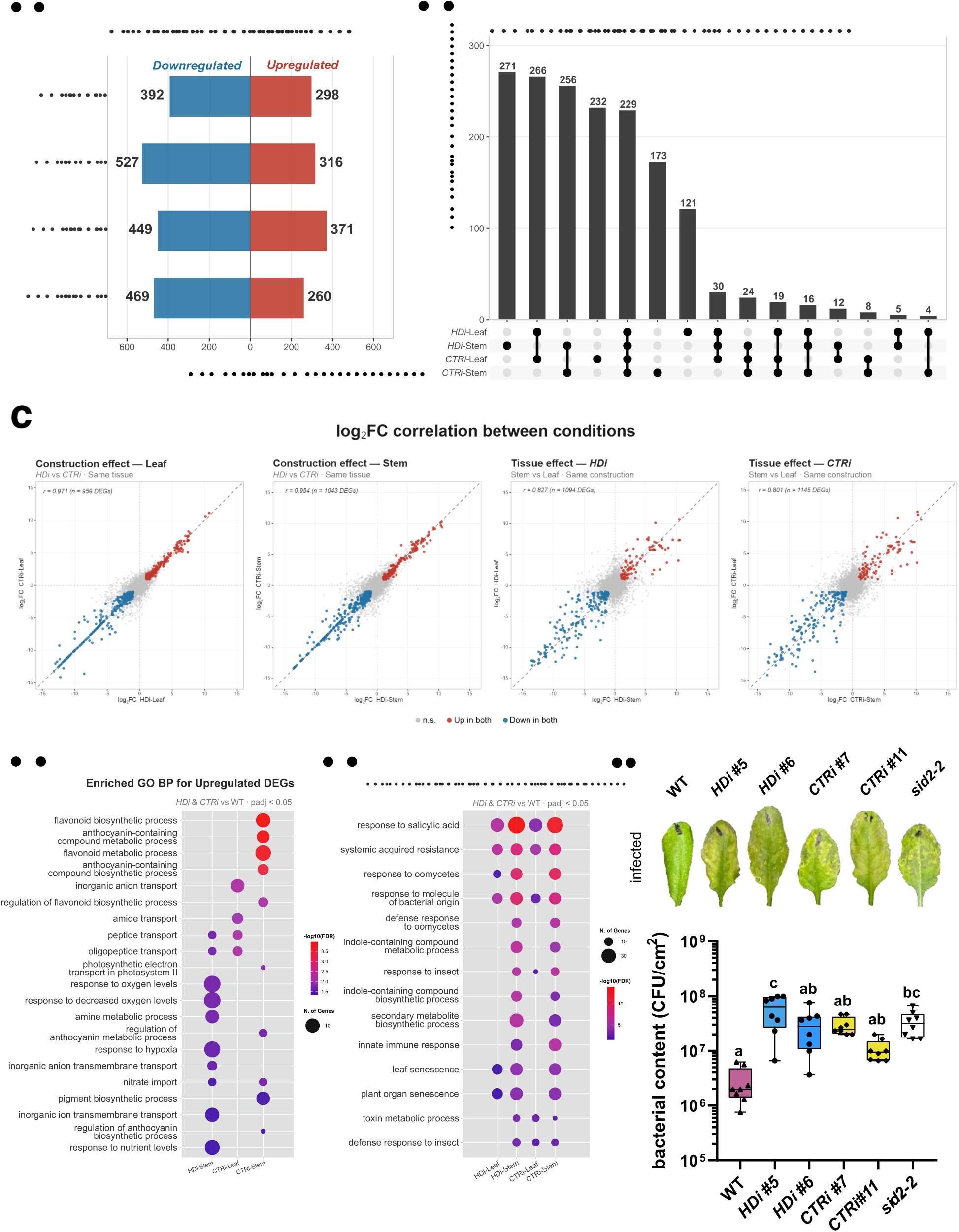
Convergent transcriptomes in plants expressing *HaHB4*-derived siRNAs reveal a suppression of biotic stress response. (a) Number of up- and downregulated genes (log_2_ fold change > 1 or < −1, *p*-adj < 0.05) in leaves and stems of 30-day-old *HDi* or *CTRi* plants relative to the WT controls. (b) Upset plot showing the overlap in differentially expressed genes (DEGs) in leaves and stems of *HDi* or *CTRi* plants. (c) Pairwise scatterplot analysis showing the highly correlated DEGs in leaves and stems of *HDi* or *CTRi* plants (compared with the WT control). Red and blue dots represent the co-up- and co-downregulated genes, respectively. (d,e) Functional enrichment analysis of genes that are significantly up- (d) or downregulated (e), as defined in (a). Point size reflects the number of genes in each category, and *p*-values are color-coded. (f) Representative images (top) and bacterial content quantification (bottom) of leaves from 28-day-old Col-0, *sid2-2*, and two independent *HDi* or *CTRi* lines, 3 days post-infection (dpi) with *Pseudomonas syringae*. Inoculation with Pseudomonas was performed by syringe infiltration, and pathogen growth is shown as CFUs cm^-2^. Data are means (±SEM) of *n*=8 independent biological replicates. Different letters indicate significant differences among means as determined using one-way ANOVA followed by Tukey’s *post-hoc* test (*P*<0.05). CFU, colony-forming units.

To identify the biological pathways affected by *HaHB4*-derived siRNAs, we performed a Gene Ontology (GO) enrichment analysis of the DEGs. Functional analysis of the upregulated genes revealed terms associated with flavonoid biosynthesis, compound transport, and response to oxygen, showing limited overlap between the two RNAi lines (Figure 4d). In stark contrast, the down-regulated DEGs displayed a consistent and significant enrichment of GO terms associated with defense responses to biotic stress in both RNAi lines, regardless of the tissue examined (Figure 4e). Based on these transcriptomic findings, we examined whether *HDi* and *CTRi* lines displayed a compromised response to biotic stress. To test this, we grew these lines under a neutral-day photoperiod (12 h light/12 h dark) for 21 days before inoculation with *Pseudomonas syringae*, to subsequently quantify bacterial proliferation in leaf tissues at 3 days post-infection. As controls, we used WT plants and the hypersusceptible *SALICYLIC ACID INDUCTION DEFICIENT 2* (*sid2-2*) mutants, which are impaired in isochorismate-mediated salicylic acid biosynthesis (Wildermuth et al., 2001). In agreement with our transcriptomic results, the *HDi* and *CTRi* plants allowed higher bacterial growth than WT controls, indicating a degree of susceptibility comparable to that of *sid2-2* mutant plants (Figure 4f). These results suggest that the expression of *HaHB4*-derived siRNAs negatively modulates the plant immune response against bacterial infection.

### The beneficial traits of plants producing *HaHB4*-derived siRNAs do not arise from a generalized accumulation of small RNAs

Given the highly consistent transcriptomic profiles between the two siRNA-expressing genotypes, we postulated that the massive accumulation of any small RNA population could non-specifically trigger these phenotypes. To test this hypothesis, we generated transgenic plants using a genetic construct to overexpress inverted repeats of a random, non-targeting sequence (*RSi*) (Caraballo et al., 2026). We systematically quantified stem width and seed yield to assess sequence specificity. In contrast to *HDi* and *CTRi* plants, several independent *RSi* plants were indistinguishable from the WT (Figure 5a,b), suggesting that the beneficial traits observed in plants expressing *HaHB4*-derived siRNAs are sequence-dependent. Consistently, we obtained similar results in plants overexpressing the *ELONGATION FACTOR-TU RECEPTOR*-associated inverted-repeat (*Ea-IR*) transposon (Mencia et al., 2025) (Figure 5c,d).

**Figure 5.**
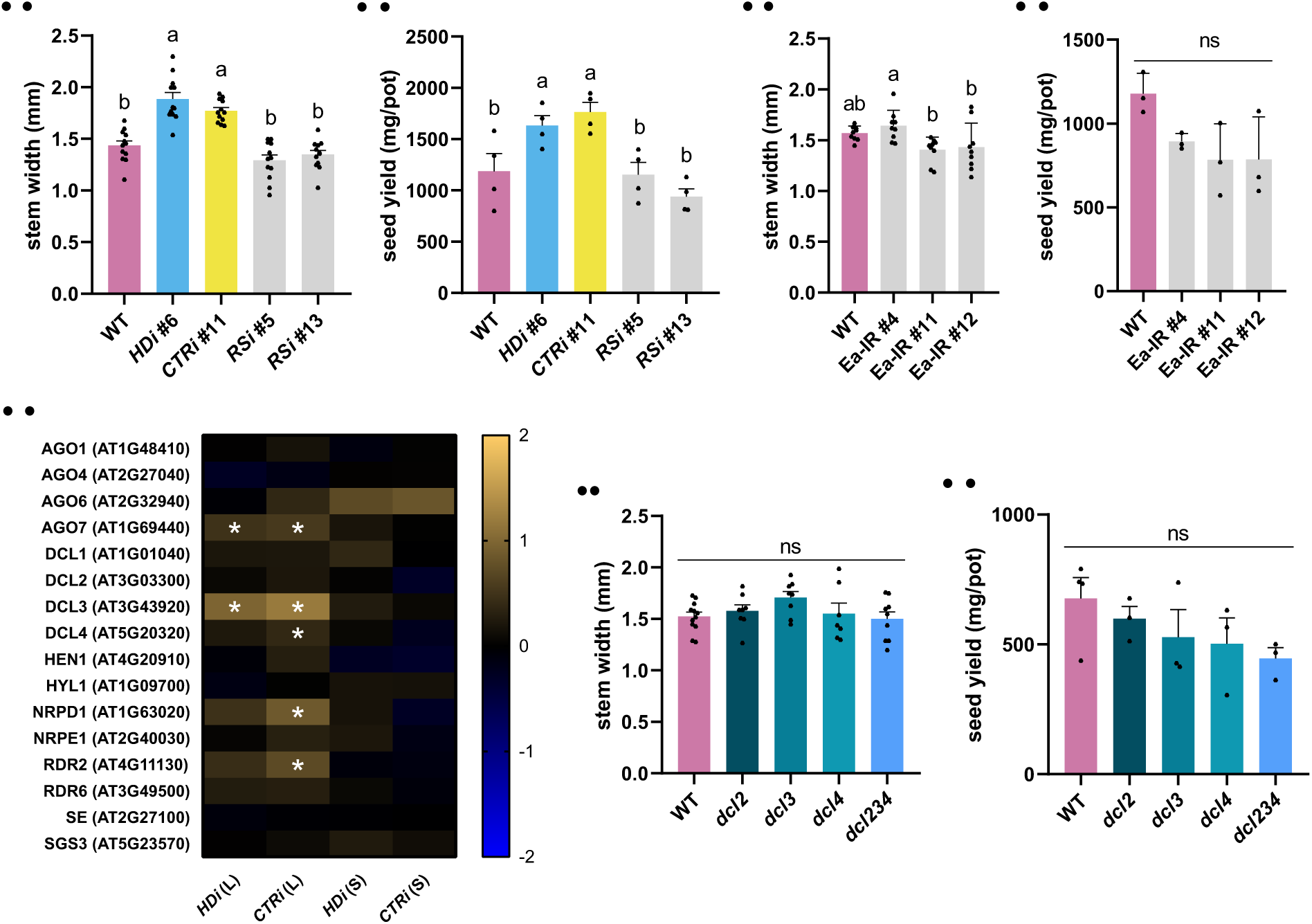
Beneficial traits induced by *HaHB4-*derived siRNAs are independent of generalized small RNA accumulation. (a) Stem width quantification of 30-day-old Col-0, *HDi*, and *CTRi* plants, and two independent lines overexpressing inverted repeats of a random, non-targeting sequence (*RSi*). (b) Seed yield evaluated at the end of the life cycle of the plants described in (a). (c) Stem width measured 30 days after sowing for Col-0 and three independent lines overexpressing the ELONGATION FACTOR-TU RECEPTOR-associated inverted-repeat (*Ea-IR*) transposon. (d) Seed yield evaluated at the end of the life cycle of the plants described in (c). (e) Heatmap of expression changes in selected small RNA pathway genes, determined by RNA-seq analysis of leaves (L) and stems (S) of 30-day-old *HDi* or *CTRi* plants. The yellow-to-blue color scale indicates log_2_(fold change) relative to the WT level for each given gene. Asterisks highlight significant differential gene expression with a *p*-adj < 0.05. (f) Stem width quantification of 30-day-old Col-0, *dcl2*, *dcl3*, *dcl4*, and *dcl2-1 dcl3-1 dcl4-2t* (*dcl234*) plants. (g) Seed yield evaluated at the end of the life cycle of the plants described in (f). For (a-g), plants were grown under long-day conditions at 22 °C, with 3 or 4 pots per genotype and three plants per pot. For (b), (d), and (g), seed yield was expressed as mg/pot. For (a)-(d) and (f)-(g), data are means (±SEM) of *n*=3-12 independent biological replicates. Different letters indicate significant differences among means as determined using one-way ANOVA followed by Tukey’s *post-hoc* test (*P*<0.05). ns, not significant.

To rule out potential feedback mechanisms involving the silencing machinery, we examined whether the expression of *HaHB4*-derived siRNAs altered the transcription of core components of the small RNA pathway. The transcriptomic analysis revealed that *HDi* and *CTRi* leaves displayed a significant upregulation of *DCL3* expression (Figure 5e). We therefore evaluated whether *dcl* loss-of-function mutants exhibited developmental phenotypes opposite to those of *HDi* and *CTRi* plants. However, the *dcl* mutants showed stem widths and seed yields similar to those of WT controls (Figure 5f,g). Altogether, these findings demonstrated that the beneficial traits of plants expressing *HaHB4*-derived siRNAs were sequence-specific and independent of generalized perturbations in the small RNA machinery.

### Plants expressing *HaHB4*-derived siRNAs exhibit partial insensitivity to salicylic acid-mediated growth repression

The growth-immunity trade-off is a highly regulated process characterized by a reciprocal suppression between development and defense, ensuring optimal energetic allocation (Huot et al., 2014). Because *HDi* and *CTRi* lines showed repression of genes related to biotic stress and increased susceptibility to pathogen infection (Figure 4), we postulated that plants expressing *HaHB4*-derived siRNAs prioritize growth over defense. To test this, we measured the primary root length in WT and the RNAi seedlings grown in the presence or absence of salicylic acid (SA). As previously reported (Pasternak et al., 2019, 2005), exogenous SA treatment significantly inhibits primary root growth in WT seedlings (Figure 6a,b). Notably, both *HDi* and *CTRi* lines were less sensitive to SA-mediated repression of root growth (Figure 6a-c), suggesting that the accumulation of *HaHB4*-derived siRNAs attenuates SA signaling pathways. Consistent with this, the RNAi lines were also less sensitive to SA-mediated inhibition of hypocotyl elongation when grown in complete darkness (Figure 6d-f). Together with the increased sensitivity to bacterial infections, these findings suggest that the expression of *HaHB4*-derived siRNAs reconfigures the balance between growth and immune responses.

**Figure 6.**
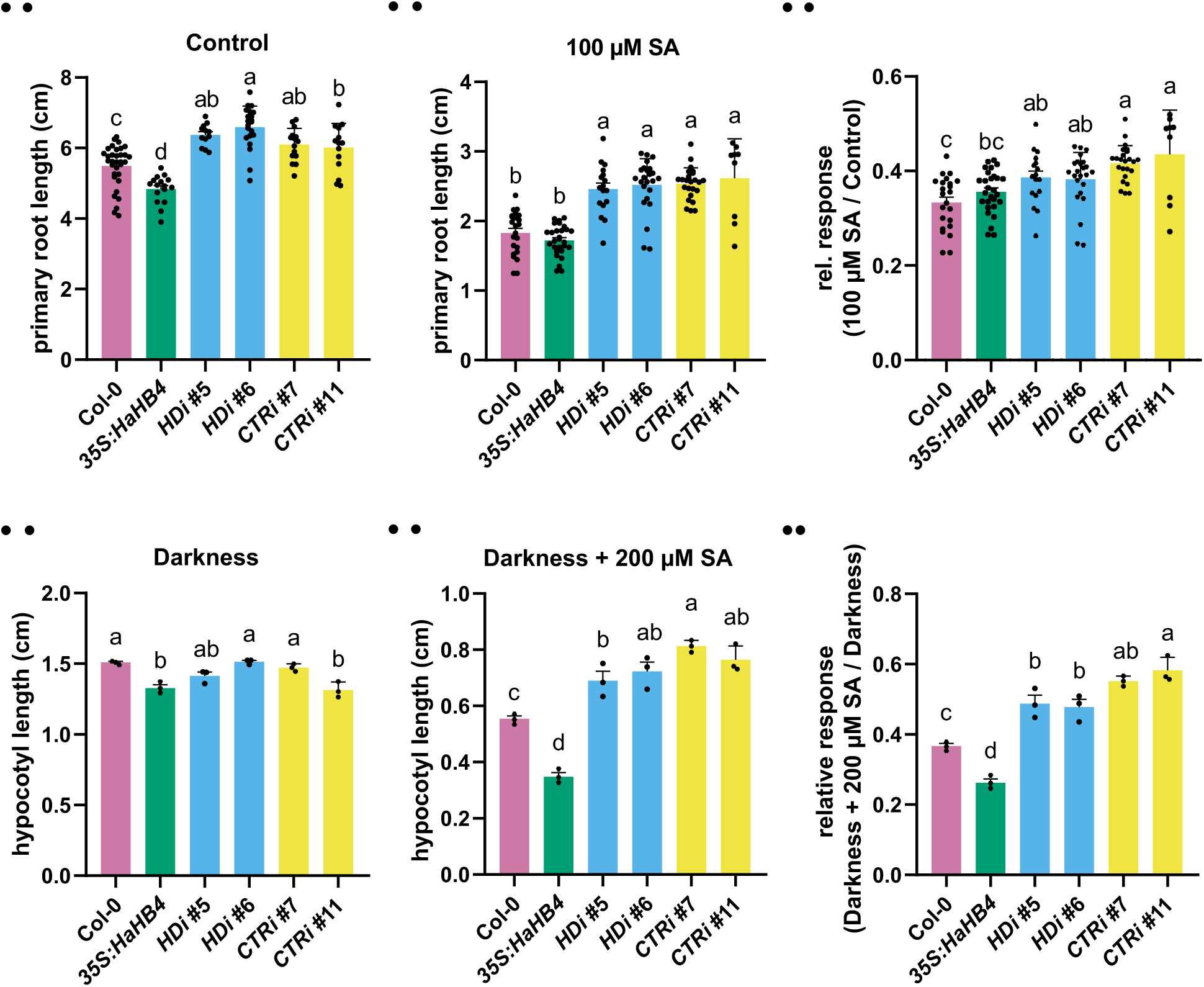
Plants producing *HaHB4*-derived siRNAs are less sensitive to salicylic acid. (a,b) Primary root length of 10-day-old Col-0, the low-expressing *HaHB4* line 30 (*35S:HaHB4*), and two independent *HDi* or *CTRi* lines grown under long-day conditions at 22 °C, in the absence (a) or presence (b) of salicylic acid (SA) (100 μM). (c) Relative root elongation response to SA for the genotypes in (a,b), calculated as the ratio of primary root length under SA treatment relative to control conditions. (d,e) Hypocotyl length of 3-day-old Col-0, *35S:HaHB4*, and two independent *HDi* or *CTRi* lines grown under continuous darkness, in the absence (d) or presence (e) of SA (200 μM). (f) Relative hypocotyl elongation response to SA for the genotypes in (d,e), calculated as the ratio of hypocotyl length under SA treatment relative to control conditions. All data are means (±SEM) of *n*=3-33 independent biological replicates. Different letters indicate significant differences among means as determined using one-way ANOVA followed by Tukey’s *post-hoc* test (*P*<0.05).

### *HaHB4*-derived siRNAs might influence endogenous Arabidopsis HD-Zip I genes, improving plant performance

Since *HaHB4* encodes a sunflower HD-Zip I transcription factor, we hypothesized that transgene-derived siRNAs might target and repress homologous Arabidopsis genes of the same subfamily. In agreement with this hypothesis, *in silico* predictions revealed that *HaHB4*-derived siRNAs target several endogenous HD-Zip I transcripts (Table S5). However, our RNA-Seq data did not show significant differences in the expression of endogenous HD-Zip I genes in *HDi* and *CTRi* lines relative to WT in either leaves or stems (Figure S4a). Because small RNAs can inhibit translation without altering target mRNA abundance (Brodersen et al., 2008; Chen, 2004), we decided to characterize T-DNA insertional mutants of candidate HD-Zip I genes predicted to be targets with known or predicted roles in shoot and stem architecture. We first evaluated the paralogs AtHB21, AtHB40, and AtHB53, which have been implicated in the repression of branch development under limiting-light conditions (González-Grandío et al., 2017). Although *athb40* single and *athb40/athb21* double mutants displayed no obvious differential phenotypes, two independent knockout alleles for *AtHB53* (*athb53-1* and *athb53-2*) exhibited a significant increase in cauline branch number while maintaining seed yield comparable to WT (Figure S4b-e). We next examined AtHB20 and AtHB3, two HD-Zip I members predicted to regulate cell wall remodeling and whose closest paralogs act as negative regulators of stem elongation (Nolan et al., 2023; Ribone et al., 2015). We found that three independent *athb20* (*athb20-1*, *athb20-2*, and *athb20-3*) lines displayed significantly longer and wider stems compared to WT (Figure S4f). Conversely, plants overexpressing *AtHB20* exhibited shorter, thinner stems than untransformed controls (Figure S4g), firmly establishing AtHB20 as a negative regulator of primary stem growth and radial expansion. Notably, neither loss- nor gain-of-function *AtHB20* lines exhibited significant differences in seed yield (Figure S4h). In contrast, loss of *AtHB3* (*athb3-1*) did not exhibit significant stem growth phenotypes under control conditions (Figure S4i). Collectively, these findings suggest that *HaHB4*-derived siRNAs might attenuate *AtHB20* and *AtHB53* expression, increasing stem width and cauline branching, respectively.

### Expression of *HaHB4*-derived siRNA confers beneficial traits in soybean

Having established that *HaHB4*-derived siRNAs enhance developmental performance in Arabidopsis, we next investigated whether their expression in crops could confer similar beneficial traits. To test this, we transformed soybean (*Glycine max* cv. Williams 82) with the *HDi* and *CTRi* constructs, generating at least two independent transgenic lines per construct to assess phenotypic outcomes. Consistent with our findings in Arabidopsis (Figure 2), *HDi* and *CTRi* transgenic soybean lines displayed enhanced growth vigor relative to WT controls under controlled environmental conditions (Figure 7a). Specifically, these siRNA-expressing lines exhibited significantly expanded total leaf area as well as increased hypocotyl and epicotyl width, similar to the traits observed in plants expressing full-length *HaHB4* (b10H) (Figure 7b-d). Furthermore, the *HDi* and *CTRi* lines showed increases in total shoot biomass (Figure 7e). Overall, these results suggest that some of the benefits conferred by the expression of *HaHB4*-derived siRNAs in Arabidopsis successfully translate to soybean, enhancing vegetative vigor during early development.

**Figure 7.**
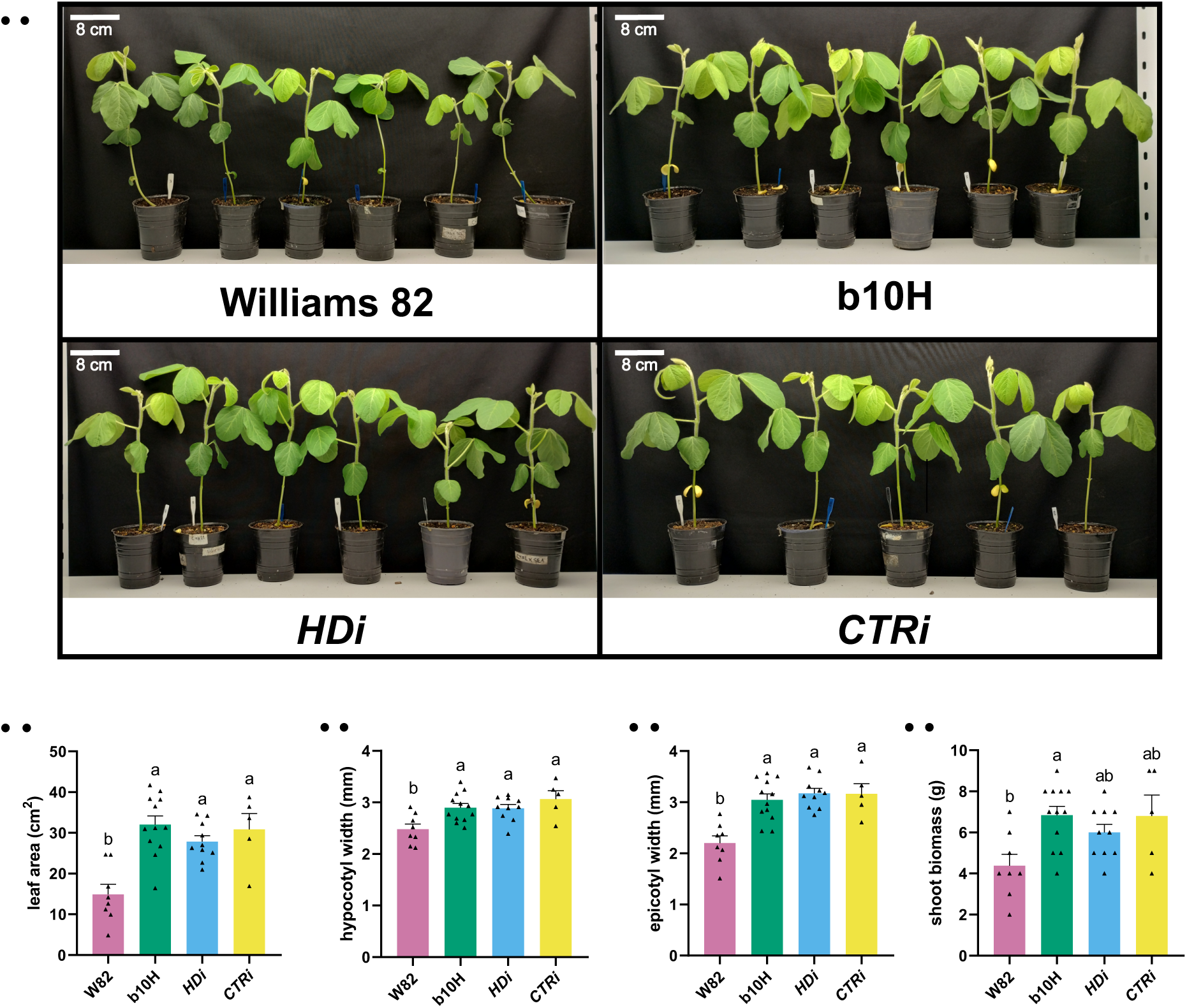
Expression of *HaHB4*-derived siRNAs enhances vegetative growth in soybean. (a) Illustrative pictures of 17-day-old WT (Williams 82, W82) and independent transgenic soybean lines expressing *HaHB4* (b10H), *HDi,* or *CTRi*. Scale bar: 8 cm. (b) Total leaf area of 13-day-old WT and transgenic plants. (c,d) Quantification of hypocotyl (c) and epicotyl (d) width in 17-day-old soybean plants. (e) Fresh weight shoot biomass of 20-day-old WT and transgenic plants. All data are means (±SEM) of *n*=5-6 independent biological replicates. Different letters indicate significant differences among means as determined using one-way ANOVA followed by Tukey’s *post-hoc* test (*P*<0.05).

## Discussion

Small RNA-based strategies have been successfully applied to improve plant performance by modulating specific endogenous targets and to enhance host immune responses (Bilir et al., 2022; Hong et al., 2024; Li et al., 2023; Shen et al., 2022; Song et al., 2018; Tao et al., 2018). In contrast, transgene-derived siRNAs are generally viewed as a mechanism that restricts excessive transgene expression (Vaucheret and Voinnet, 2024). Whether such siRNAs can acquire regulatory activity toward endogenous pathways, rather than merely promoting silencing of the transgene, remains largely unexplored. Here, we reveal an unexpected outcome of transgene silencing, since small RNAs derived from a heterologous transcription factor can generate a distinct and beneficial developmental program in Arabidopsis and soybean.

Constitutive expression of *HaHB4* in Arabidopsis has previously been associated with compact rosettes, rounded leaves, shortened petioles, and delayed flowering (Dezar et al., 2005). We found these characteristics only in a fraction of independent *35S:HaHB4* transformants in WT, whereas their frequency increased markedly in backgrounds defective in small RNA biogenesis or amplification (Figure 1). Together with previous reports of increased expression of Arabidopsis HD-Zip I genes in small-RNA mutants (Miguel et al., 2020; Ribone et al., 2017; Romani et al., 2016), the correlation between *HaHB4* transcript abundance and phenotypic severity supports the notion that *HaHB4* is subjected to RNA-silencing mechanisms in Arabidopsis.

In contrast to *35S:HaHB4* plants, expression of either of the two independent hairpin constructs derived from distinct, non-overlapping segments of the *HaHB4* coding sequence promoted vegetative growth and reproductive performance. Indeed, the *HDi* and *CTRi* plants exhibited increased hypocotyl and leaf growth, enhanced root development, wider stems, increased cauline branching, and higher seed yield (Figure 2). Remarkably, the phenotypes of both lines were abolished in the *dcl234* background (Figure 2o,p), consistent with the requirement of DCL-mediated siRNA biogenesis for hairpin RNA-induced silencing phenotypes (Fusaro et al., 2006; Mlotshwa et al., 2008). The enhanced reproductive performance of plants expressing *HaHB4*-derived siRNAs was maintained under water-limiting conditions (Figure 3), indicating that the reproductive advantage of the RNAi plants was not restricted to optimal growth conditions. However, this should not be interpreted as evidence of increased drought tolerance, but rather as an enhanced capacity to maintain reproductive output under the conditions tested. Although *HaHB4* expression enhances crop seed yield under water-limiting conditions (González et al., 2019; Ribichich et al., 2020), its overexpression in Arabidopsis produced no significant difference relative to WT plants under our experimental conditions (Figure 3). While seed yield following drought treatment had not been previously evaluated in Arabidopsis, this discrepancy may stem from differences in expression systems. For instance, *HaHB4* is driven by its native promoter in soybean, whereas it is expressed under the constitutive *35S* promoter in Arabidopsis or by the *UBI1* promoter in wheat (Dezar et al., 2005; González et al., 2019; Ribichich et al., 2020).

The similarity between the *HDi*/*CTRi* plants and the phenotype reported following RNAi-mediated downregulation of *GhADF1* (*ACTIN-DEPOLYMERIZING FACTOR 1*) in cotton (Qin et al., 2022) raises the possibility of unintended off-target effects. However, *AtADF1* was not differentially expressed in our RNA-Seq datasets (Tables S1–S4), arguing against this gene as a major target. Another possibility is that transgene-derived double-stranded RNAs or siRNAs perturb endogenous small-RNA regulatory pathways. Small-RNA pathways are important for normal reproductive development, and mutations affecting their components can strongly impair fertility and seed production (Oliver et al., 2017; Peragine et al., 2004; Wei et al., 2020). Nevertheless, plants expressing unrelated inverted-repeat sequences did not reproduce the increased stem width or seed yield (Figure 5), suggesting that the effects of the *HaHB4*-derived siRNAs are specific and sequence-dependent.

Previous studies have linked increased stem diameter and vascular development with enhanced seed yield in Arabidopsis (Cabello et al., 2016; Cabello and Chan, 2019; Raminger et al., 2025, 2023), strongly suggesting that modifications in stem anatomy contribute to reproductive performance. In agreement, tomato plants exhibiting wider stems provoked by mechanical treatment also showed enhanced fruit yield (Castro-Estrada et al., 2024). Accordingly, the expression of *HaHB4*-derived siRNAs promoted increased stem and pith areas and higher seed yield without increasing vascular bundle number (Figures 2 and S1). Notably, auxin transporter mutants showed increased stem diameter following mechanical treatment without increasing vascular bundle number or seed yield (Cabello and Chan, 2019), indicating that stem expansion alone is unlikely to account for the increased reproductive output. The close association between seed yield, cauline branch number, and silique number in the RNAi lines (Figure 2k,l) instead suggests that stem expansion and increased reproductive output may represent parallel consequences of the siRNA-induced developmental program. Alternatively, wider stems could contribute to the increased sink demand associated with a higher silique number by facilitating photoassimilate transport. Indeed, the coordination of carbohydrate partitioning between source and reproductive sink tissues is a major determinant of seed yield (Pegler et al., 2023; Smith et al., 2018).

Despite being derived from distinct regions of *HaHB4*, the *HDi* and *CTRi* constructs produced remarkably similar phenotypes and largely overlapping transcriptional changes (Figures 2 and 4). Notably, most repressed genes were associated with biotic stress responses, consistent with the increased susceptibility to bacterial infection observed in these plants (Figure 4e,f). This phenotype is similar to that of *HaHB4*-overexpressing Arabidopsis plants, which display increased susceptibility to Pseudomonas and reduced SA levels (Manavella et al., 2008b). Moreover, *HDi* and *CTRi* plants showed reduced sensitivity to exogenous SA during development (Figure 6). These observations suggest that the RNAi lines modify the balance between growth and immunity, contributing to their enhanced growth and reproductive output (Figure 2). Consistent with this, SA-deficient Arabidopsis plants exhibit increased biomass and seed production (Abreu and Munne-Bosch, 2009).

The similar transcriptomes and phenotypes generated by the two non-overlapping hairpins suggest that distinct *HaHB4*-derived siRNA populations may converge on common endogenous regulatory pathways, even without targeting identical individual transcripts. Because siRNAs can silence off-target transcripts through partial sequence complementarity (Ossowski et al., 2008), another possibility is that *HaHB4*-derived siRNAs may function as a multi-target module acting on endogenous genes with high sequence similarity to *HaHB4*. Supporting the latter hypothesis, loss-of-function of endogenous HD-Zip I genes produces developmental traits that overlap with those altered in *HDi* and *CTRi* plants. AtHB21, AtHB40, and AtHB53 have been implicated in branching repression under light-limiting conditions (González-Grandío et al., 2017), and we found that AtHB53 inhibits cauline branching (Figure S4d). We also identified AtHB20 as a negative regulator of primary stem elongation and radial expansion (Figure S4f,g). Neither *athb20* nor *athb53* single mutants showed enhanced seed yield despite displaying specific beneficial traits (Figure S4). We cannot exclude the possibility that *HaHB4*-derived siRNAs modulate the expression of additional genes. For example, loss of the related HD-Zip I gene *AtHB5* increases stem width and seed yield (Raminger et al., 2023). Together, these observations indicate that *HaHB4*-derived siRNAs simultaneously affect multiple endogenous HD-Zip I regulatory pathways, generating a composite phenotype that cannot be fully recapitulated by mutating a single gene. Given that the *HDi* and *CTRi* lines exhibited reduced sensitivity to SA (Figures 4 and 6), determining whether single or multiple HD-Zip I family members govern these biotic responses represents an important direction for future research.

A limitation of this model is that neither *AtHB20* nor *AtHB53* transcript abundance was significantly altered in the bulk RNA-Seq datasets (Figure S4a). This does not exclude their regulation at the translational level or in specific cell types. Indeed, small RNAs can repress gene expression through translation inhibition in addition to transcript degradation (Borges and Martienssen, 2015). For example, an endogenous 24-nt small RNA generated by RDR2 and DCL3 regulates the HD-Zip IV gene *OCL1* (*Outer Cell Layer 1*) primarily via translational repression (Klein Cosson et al., 2015). Furthermore, *AtHB53* is expressed specifically in young axillary meristems and at the base of leaf primordia in buds (González-Grandío et al., 2017), and transcriptional changes could be diluted in bulk tissue. Future experiments will be required to determine whether *AtHB20*, *AtHB53*, or other endogenous transcripts are directly regulated by the *HaHB4*-derived siRNAs.

Importantly, several differential traits presented by *HDi* and *CTRi* Arabidopsis plants were also observed in soybeans (Figure 7). Even though further studies and field trials will be necessary to reach more supported conclusions, this appears to be an example of successful translation from the model plant Arabidopsis to a crop. As previously described (Uauy et al., 2025), such success was not routine in the past.

Together, our findings demonstrate that transgene-derived siRNAs can act as biologically active molecules able to modulate endogenous gene networks to drive complex phenotypic outputs, rather than functioning solely as a barrier to transgene expression. Although the precise targets and mechanisms remain to be determined, our results suggest that heterologous gene-derived siRNAs can contribute to regulatory diversity and may provide an additional route for engineering complex agronomic traits, highlighting their potential as powerful biotechnological tools.

## Materials and Methods

### Plant material

Wild-type and other lines of plants used in this study were *Arabidopsis thaliana* L. Heynh. ecotype Columbia (Col-0). The mutant seeds *athb53-1* (GK-085A06-011845; CS408070), *athb53-2* (GK-312E06-015817; CS429910), *dcl2-1* (SALK_064627), *dcl3-1* (SALK_005512), *dcl4-2* (GABI_160G05), *dcl2-1 dcl3-1 dcl4-2t* (*dcl234*; CS16391), *rdr2-1* (SAIL_1277_H08; CS879934), and *rdr6-12* (CS24286) in the Col-0 ecotype background were obtained from the Arabidopsis Biological Resource Center (ABRC; Columbus, OH, USA; http://www.arabidopsis.org). The mutant plants *athb20-1* (SALK_064218), *athb20-2* (SALK_204744), *athb20-3* (SAIL_300_B06), *athb40-1*, *athb40-2*, and *athb40-2 athb21*, and plants overexpressing *AtHB20* (AT20.1, AT20.2, and AT20.3) or *HaHB4* (lines *35S::Hahb4*-6 and *35S:Hahb4*-30) were previously described (Cabello et al., 2007; Dezar et al., 2005; González-Grandío et al., 2017; Mora et al., 2022; Murguía et al., 2026b). Soybean plants (Williams 82 genotype) expressing the sunflower *HaHB4* gene were previously described (Ribichich et al., 2020).

### Plant growth conditions and treatments

Arabidopsis plants were grown on soil in a 21-23 °C growth chamber under a long-day photoperiod (16 h light, 110 μmol m^−2^ s^−1^/8 h dark). For *in vitro* growth, seeds were surface-sterilized with 70% EtOH followed by 10% sodium hypochlorite, thoroughly rinsed with sterile water, and stratified at 4 °C for 3 days to ensure uniform germination. Unless stated otherwise, seedlings were grown in Petri dishes containing half-strength Murashige and Skoog (½MS) basal medium (Murashige and Skoog, 1962) supplemented with vitamins (PhytoTechnology Laboratories) and 0.9% agar, at 21-23 °C under long-day conditions (16 h light, 110 μmol m^−2^ s^−1^/8 h dark). For hypocotyl elongation, seeds were grown in ½MS medium containing 200 µM salicylic acid at 22 °C under complete darkness. Soybean plants were grown in 0.5 l pots (one plant per pot) in a 28 °C growth chamber under a long-day photoperiod (16 h light, 250 μmol m^−2^ s^−1^/8 h dark).

### Genetic constructs

Cloning was performed using standard molecular biology procedures. For the *RNAi-HB4HD* (*HDi*) construct, a 227-bp fragment corresponding to the HaHB4 homeodomain was amplified by PCR using the oligonucleotides RNAi HDS F and RNAi HDS R (sense orientation) or RNAi HDAS F and RNAi HDAS R (antisense orientation). The sense and antisense fragments were cloned into the *pHANNIBAL* vector using the *Xho*I/*Eco*RI and *Hind*III/*Bam*HI restriction sites, respectively. The resulting RNAi cassette was subsequently excised and subcloned into the *Sac*I/*Pst*I sites of the binary vector *pTF101.3*. For the *RNAi-HB4CTR* (*CTRi*) construct, a 309-bp fragment corresponding to the HAHB4 leucine zipper and carboxy-terminal region was amplified by PCR using the oligonucleotides RNAi CTRS F and RNAi CTRS R (sense orientation) or RNAi CTRAS F and RNAi CTRAS R (antisense orientation). The sense and antisense fragments were cloned into the *pHANNIBAL* vector using the *Xho*I/*Eco*RI and *Hind*III/*Bam*HI restriction sites, respectively. The resulting RNAi cassette was subsequently excised and subcloned into the *Sac*I/*Pst*I sites of the binary vector *pTF101.3*. The new clones were checked for correctness by DNA sequence analysis (Macrogen, Korea). A complete list of plasmids used in this study is provided in Table S6. Plasmid maps and DNA sequences are available upon request.

### Plant transformation

Arabidopsis plants were stably transformed via floral dip using *Agrobacterium tumefaciens* LBA4404 harboring the specific constructs (Clough and Bent, 1998). Transgenic plants were selected on ½ MS medium supplemented with the appropriate selective agent (ammonium glufosinate 1 ml l^-1^, or kanamycin 25 mg l^-1^). Seeds were surface-sterilized, stratified for 3 days at 4 °C, and then germinated in a growth chamber at 21-23 °C. When ammonium glufosinate was used, selection was performed directly in soil. For each construct, three to four independent positive lines confirmed by genotyping (primer sequences listed in Table S7) were propagated, and homozygous T3 or T4 plants carrying a single insertion, as determined by Mendelian segregation, were used for further analyses.

Soybean transgenic lines were generated using the *RNAi-HB4HD* (*HDi*) and *RNAi-HB4CTR* constructs, an Agrobacterium-mediated protocol, and the wild-type (WT) cultivar Williams 82 (W82) according to the methods described by Somers et al. (2003). Transgenic lines were selected using ammonium glufosinate. T1 seeds were obtained for four independent lines for each construct. Seed multiplication was conducted in a greenhouse. T1 individuals derived from each genotype were sampled for a segregation test using PCR. Genotypes derived from selfings of individuals (3:1 segregation in T1) were sown and analyzed by PCR to identify homozygous lines, as indicated by the absence of negative segregants among the sampled progeny. Seed augmentation (T3 seed) of single-copy homozygous lines was carried out in a greenhouse.

### Measurement of hypocotyl length

Hypocotyl length was quantified as the distance from the most basal root hair to the “V” junction formed by the cotyledons, as previously reported (Capella et al., 2015). All measurements were performed using Fiji/ImageJ software (Schindelin et al., 2012). Each biological replicate was calculated as the mean hypocotyl length of at least 15 seedlings cultivated on the same plate. Graphical data represent the mean values obtained from 3-4 independent plates per treatment and/or genotype.

### Plant phenotyping

Plant architectural traits were quantified either manually or with the aid of a ruler or gauge. For each genotype and treatment, 4-8 Arabidopsis plants were analyzed, and all experiments were independently repeated three times with comparable results. Stem diameter was measured at the first internode of wild-type or transgenic Arabidopsis plants with a stem height of 10 cm. For seed yield determination, plants were grown at a density of three individuals per pot and harvested at maturity. Yield was expressed as grams of seed per pot. For root length measurements, surface-sterilized seeds were sown 1 cm from the top of square Petri dishes (12 × 12 cm) containing ½ MS medium containing 100 µM salicylic acid and grown vertically under long-day conditions for the indicated period of time. Seedlings were photographed, and root length was quantified using Fiji/ImageJ (Schindelin et al., 2012). For total exposed leaf area, 15-day-old plants were photographed, and their areas were analyzed using the same image-processing software. For soybean, six plants were analyzed per experiment (one representative line per transgenic event) grown at a density of one plant per pot, and all experiments were independently repeated three times with consistent results. In 17-day-old wild-type and transgenic plants, hypocotyl and epicotyl diameters were measured 2 cm above the soil level and immediately above the cotyledonary node, respectively. To determine shoot biomass, the weight of tissues excised above the cotyledons was measured in 20-day-old plants. Additionally, leaves of 13-day-old wild-type and transgenic plants were photographed, and total leaf area was quantified using Fiji/ImageJ (Schindelin et al., 2012).

### Histological analysis of stem cross-sections

Arabidopsis stems were fixed in 70% ethanol for 48 h. Freehand cross-sections were prepared under a stereo microscope (Leica EZ4) on slides (25.4 × 76.2 mm) using commercial razor blades and stained with Astra blue-Safranin (Arend et al., 2008), as previously described (Castro-Estrada et al., 2024). Microscopic optical visualization (Eclipse E200, Carl Zeiss, Axiostar Plus, Göttingen, Germany) using a 10× lens was conducted to determine the stem and pith area (mm^2^) and the number of vascular bundles. Images were analyzed using Fiji/ImageJ (Schindelin et al., 2012).

### RNA isolation and analysis

Transcript levels were analyzed by RT-qPCR using RNA extracted from rosette leaves of soil-grown Arabidopsis plants. For each genotype, 3-4 biological replicates were included per experiment. Total RNA was purified from seedlings using Trizol^®^ reagent (Invitrogen) according to the manufacturer’s instructions. One μg of RNA was reverse-transcribed using oligo(dT)_18_ and M-MLV reverse transcriptase II (Promega). Quantitative real-time PCR (qPCR) assays were performed using a StepOne equipment (Applied Biosystems); each reaction contained a final volume of 20 μl that included 2 μl of SYBR Green (4×), 8 pmol of each primer, 2 mM MgCl_2_, 10 μl of a 1/20 dilution of the RT reaction, and 0.1 μl of Taq Polymerase (Invitrogen), using standard protocols (40 to 45 cycles, 60 °C annealing). Fluorescence was quantified at 72 °C. Specific primers for each gene were designed and are listed in Table S8. The expression levels were normalized using *ACTIN2/8* (AT3G18780/AT1G49240) as a reference gene, and quantification was carried out using the ΔΔCt method (Pfaffl, 2001) relative to Col-0 seedlings grown at 22 °C.

### RNA-Seq analysis

For RNA-seq analysis, total RNA was extracted from rosette leaves or stems of soil-grown Arabidopsis plants measuring 5-10 cm in height (approximately 30-day-old plants), including Col-0 wild-type or Col-0 lines transformed with the *HDi* or *CTRi* constructs. Three biological replicates per genotype were sequenced by Novogene (Sacramento, CA, USA; Project number: H202SC24128592). Libraries were sequenced on the Illumina NovaSeq X-Plus platform, generating 150-bp paired-end reads with a minimum depth of 12 million reads per sample. RNA-seq reads were analyzed on the Galaxy platform (Abueg et al., 2024). Briefly, the quality-filtered reads were aligned to the *Arabidopsis thaliana* reference genome (TAIR10) using RNA STAR (Galaxy Version 2.7.11a+galaxy1), guided by the gene and exon annotation from Araport V11. Differential expression analysis was performed using DESeq2 (Version 1.42.1) in RStudio. Functional enrichment analysis was performed using clusterProfiler 4.10.1 package (Wu et al., 2021) in RStudio. UpSet plots were generated using the packages in RStudio. RNA-seq data were deposited in the Gene Expression Omnibus (GEO) database.

### In silico prediction of siRNA targets

To identify potential target genes, the complete sequences of the *HDi* and *CTRi* constructs were in silico cleaved into all possible 21- and 24-nucleotide fragments. The generated sequence libraries were subsequently screened against the HD-Zip I gene family using the psRNATarget server (Dai et al., 2018) with default parameters, using Expectation 5.0 as a threshold.

### Statistics and reproducibility

All experiments were performed at least three times with similar outcomes, and each figure panel displays representative results from these repetitions. Analyses of variance were performed, and pairwise differences were evaluated with Tukey or Fisher’s *post hoc* test using R statistical language (R Development Core Team, 2008); different groups are marked with letters at the 0.05 significance level. For all error bars, data are mean ± S.E.M. *P* values were generated using two-tailed Student’s *t*-tests; N/S, *P* ≥ 0.05, *\*P* < 0.05, *\*\*P* < 0.01, *\*\*\*P* < 0.001.

## Supplementary data

Table S1: Differentially expressed genes in *HDi* lines in leaves

Table S2: Differentially expressed genes in *HDi* lines in stems

Table S3: Differentially expressed genes in *CTRi* lines in leaves

Table S4: Differentially expressed genes in *CTRi* lines in stems

Table S5: List of predicted siRNA targets

Table S6: Set of primers used in this study for cloning

Table S7: Set of primers used in this study for genotyping

Table S8: Set of primers used in this study for RT-qPCR experiments

## Acknowledgments

We are very grateful for the technical assistance provided by Dr. Mabel Campi, Mr. Manuel Franco, and Dr. Rafael Ambrosio for obtaining and multiplying transgenic soybean plants. We also thank Dr. Agustin Arce and Dr. Pablo Manavella for helpful discussions.

## Author contributions

M.C. and R.L.C. conceived the study. G.J.V., M.C., and R.L.C. designed experiments. M.L.B. and D.A.C. performed the biotic stress experiments. M.C. and G.J.V. analyzed the RNA-Seq data. J.E.G., E.W., and RLC performed the experiments with soybean. J.P.M. performed the analysis of AtHB3 and AtHB20 plants. L.C. generated the construct and the plants for random RNAi. G.J.V. performed all other experiments. G.J.V., M.C., and R.L.C. analyzed the project. M.C. and R.L.C. supervised the project. E.W., D.A.C., M.C., and R.L.C. acquired funding. M.C. and R.L.C. conceived and wrote the manuscript. All the authors contributed to the data analysis and discussion, and reviewed, revised, and approved the manuscript.

## Data availability

All data supporting the findings of this study, including supplementary materials, are available from the corresponding author upon request.

## Supplementary Figure legends

**Figure S1. *HDi* and *CTRi* lines exhibit stem and pith expansion without altering vascular bundle number** (a-d) Quantification of stem (a) or pith area (b), the ratio between stem and pith area (c), and number of vascular bundles (d) of 30-day-old Col-0, the low-expressing *HaHB4* line 30 (*35S:HaHB4*), and two independent Col-0 lines transformed with either the *HDi* or *CTRi* constructs. Plants were grown under long-day conditions at 22 °C, with 3 or 4 pots per genotype and three plants per pot. All data are means (±SEM) of *n*=4 independent biological replicates. Different letters indicate significant differences among means as determined using one-way ANOVA followed by Fisher’s *post-hoc* test (*P*<0.05). ns, not significant.

**Figure S2**. **Expression of *HaHB4*-derived siRNA enhances plant performance under short-day conditions** (a,b) Representative pictures (a) and quantification of hypocotyl length (b) from 6-day-old Col-0, the low-expressing *HaHB4* line 30 (*35S:HaHB4*), and two independent Col-0 lines transformed with either the *HDi* or *CTRi* constructs. (c,d) Quantification of stem width (c) and number of cauline branches (d) of 56-day-old plants described in (a). (e) Seed yield evaluated at the end of the life cycle of the plants described in (a), expressed as mg/pot. (f) Representative images of the plants described in (a). Scale bar: 4 cm. Plants were grown under short-day conditions at 22 °C, with 3 or 4 pots per genotype and three plants per pot. For (b-e), data are means (±SEM) of *n*=3-15 independent biological replicates. Different letters indicate significant differences among means as determined using one-way ANOVA followed by Tukey’s *post-hoc* test (*P*<0.05).

**Figure S3**. **Global leaf transcriptomes of *HDi* and *CTRi* lines exhibit minimal overlap with *35S:HaHB4* plants** (a) Number of up- and downregulated genes (log_2_ fold change > 1 or < −1, *p*-adj < 0.05) in leaves of 30-day-old *HDi* or *CTRi* plants relative to the WT controls (this work) and of *HaHB4*-overexpressing lines compared to WT under normal watering conditions (Manavella et al., 2006). (b) Upset plot showing the overlap in differentially expressed genes (DEGs) in leaves of *HDi*, *CTRi,* or *35S:HaHB4* plants.

**Figure S4**. ***AtHB20* and *AtHB53* loss-of-function mutants recapitulate distinct traits of plants expressing HaHB4-derived siRNA** (a) Heatmap of expression changes in all HD-Zip I genes, determined by RNA-seq analysis of leaves (L) and stems (S) of 30-day-old *HDi* or *CTRi* plants. The yellow-to-blue color scale indicates log_2_(fold change) relative to the WT level for each given gene. (b) Number of cauline branches of 40-day-old Col-0, *HDi*, *CTRi*, *athb40-1*, *athb40-2*, and *athb40-2 athb21* plants. (c) Seed yield evaluated at the end of the life cycle of the plants described in (b). (d) Number of cauline branches of 40-day-old Col-0, *HDi*, *athb53-1*, and *athb53-2* plants. (e) Seed yield evaluated at the end of the life cycle of the plants described in (d). (f) Stem length (left) and width (right) in Col-0 and three independent *athb20* mutant plants (*athb20-1*, *athb20-2*, and *athb20-3*) grown for 36 days. (g) Stem length (left) and width (right) in Col-0 and three independent *AtHB20*-overexpressing lines (*AT20.1*, *AT20.2*, and *AT20.3*) grown for 36 days. (h) Seed yield evaluated at the end of the life cycle of the plants described in (f) and (g). (i) Stem length (left) and width (right) in 36-day-old Col-0 and *athb3* plants. All plants were grown under long-day conditions at 22 °C, with 3 or 4 pots per genotype and three plants per pot. For (c), (e), and (h), seed yield was expressed as mg/pot. All data are means (±SEM) of *n*=3-45 independent biological replicates. Different letters indicate significant differences among means as determined using one-way ANOVA followed by Tukey’s *post-hoc* test (*P*<0.05). ns, not significant.

