## Supplementary Data for "Protein-independent regulation by transgene-derived small interfering RNAs rewires endogenous regulatory networks to enhance plant growth, architecture, and drought performance"

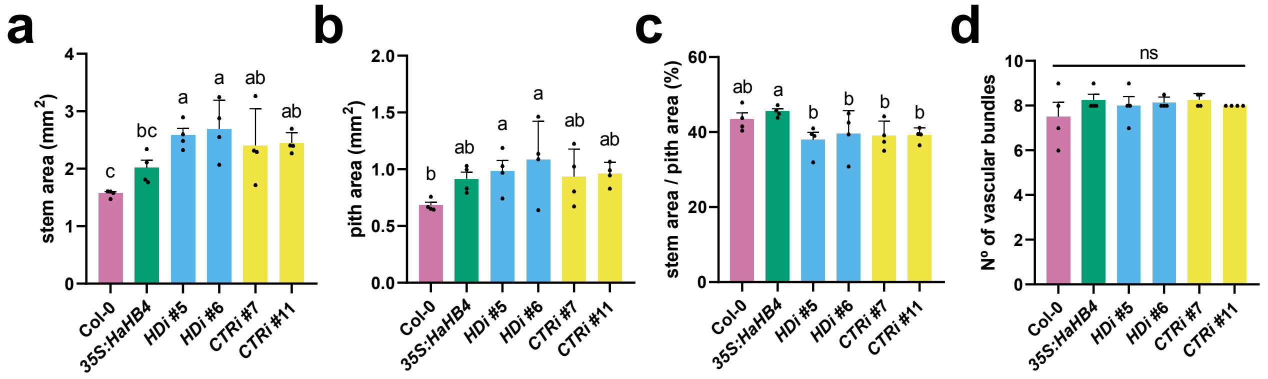


**Figure S1. *HDi* and *CTRi* lines exhibit stem and pith expansion without altering vascular bundle number**

(a-d) Quantification of stem (a) or pith area (b), the ratio between stem and pith area (c), and number of vascular bundles (d) of 30-day-old Col-0, the low-expressing *HaHB4* line 30 (*35S:HaHB4*), and two independent Col-0 lines transformed with either the *HDi* or *CTRi* constructs. Plants were grown under long-day conditions at 22 °C, with 3 or 4 pots per genotype and three plants per pot. All data are means (±SEM) of $n$=4 independent biological replicates. Different letters indicate significant differences among means as determined using one-way ANOVA followed by Fisher's *post-hoc* test ($P$<0.05). ns, not significant.

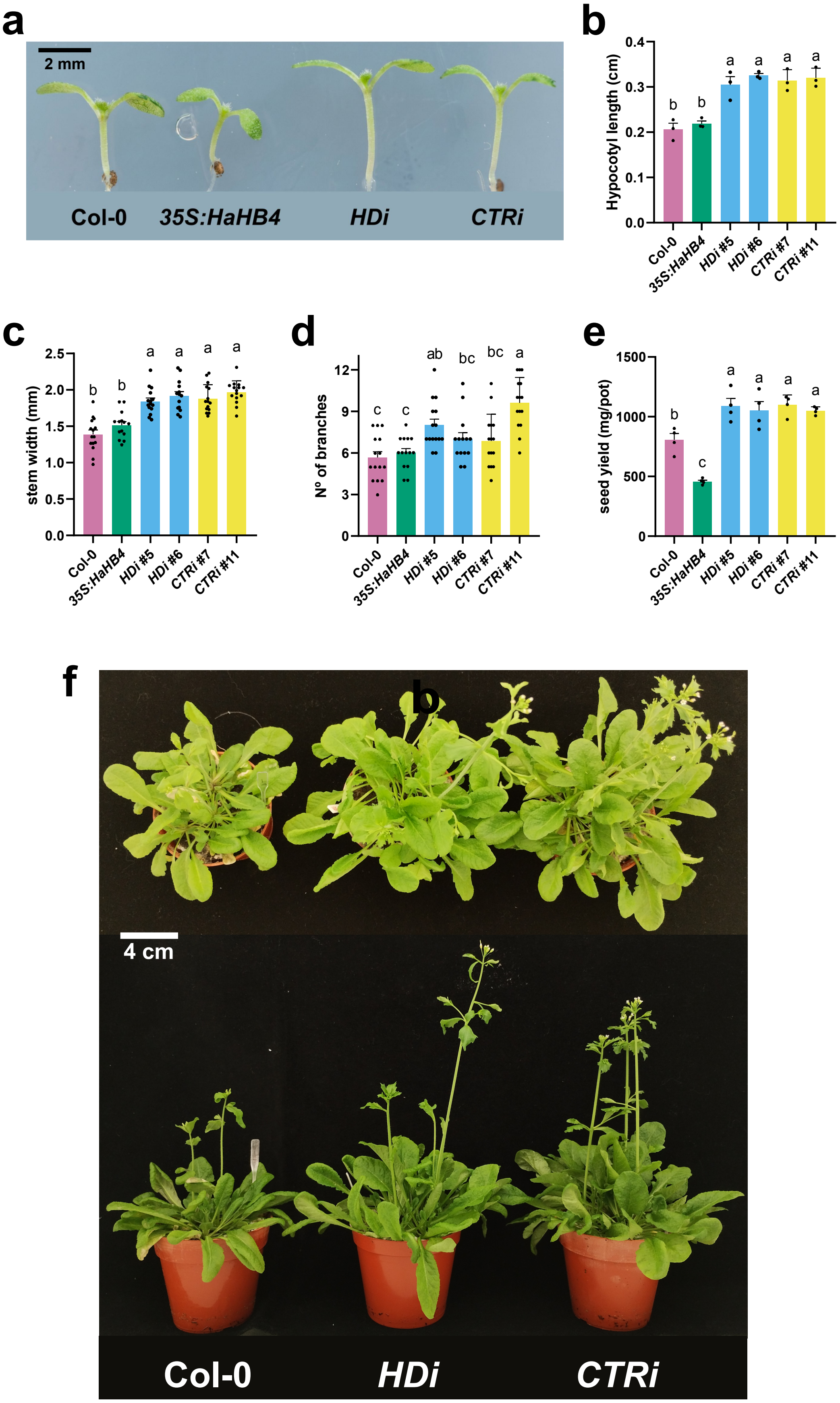


**Figure S2. Expression of *HaHB4*-derived siRNA enhances plant performance under short-day conditions**
(a,b) Representative pictures (a) and quantification of hypocotyl length (b) from 6-day-old Col-0, the low-expressing *HaHB4* line 30 (*35S:HaHB4*), and two independent Col-0 lines transformed with either the *HDi* or *CTRi* constructs. (c,d) Quantification of stem width (c) and number of cauline branches (d) of 56-day-old plants described in (a). (e) Seed yield evaluated at the end of the life cycle of the plants described in (a), expressed as mg/pot. (f) Representative images of the plants described in (a). Scale bar: 4 cm. Plants were grown under short-day conditions at 22 °C, with 3 or 4 pots per genotype and three plants per pot. For (b-e), data are means (±SEM) of $n$=3-15 independent biological replicates. Different letters indicate significant differences among means as determined using one-way ANOVA followed by Tukey's *post-hoc* test ($P$<0.05).

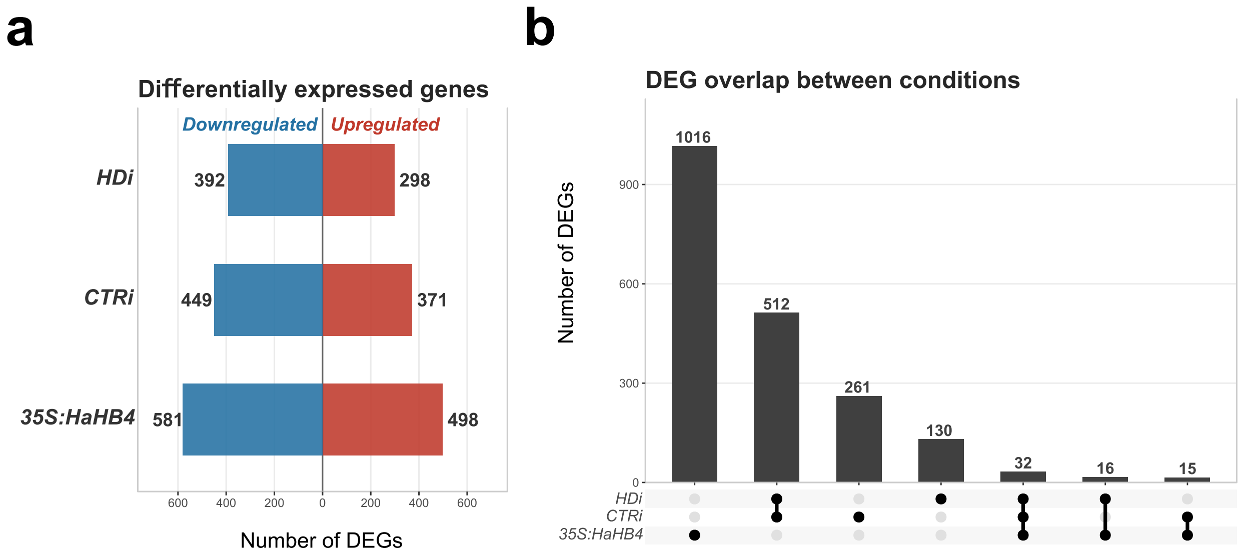


**Figure S3. Global leaf transcriptomes of HDi and CTRi lines exhibit minimal overlap with 35S:HaHB4 plants**
(a) Number of up- and downregulated genes ($log_2$ fold change > 1 or < -1, *p*-adj < 0.05) in leaves of 30-day-old *HDi* or *CTRi* plants relative to the WT controls (this work) and of *HaHB4*-overexpressing lines compared to WT under normal watering conditions (Manavella et al., 2006). (b) Upset plot showing the overlap in differentially expressed genes (DEGs) in leaves of *HDi*, *CTRi*, or *35S:HaHB4* plants.

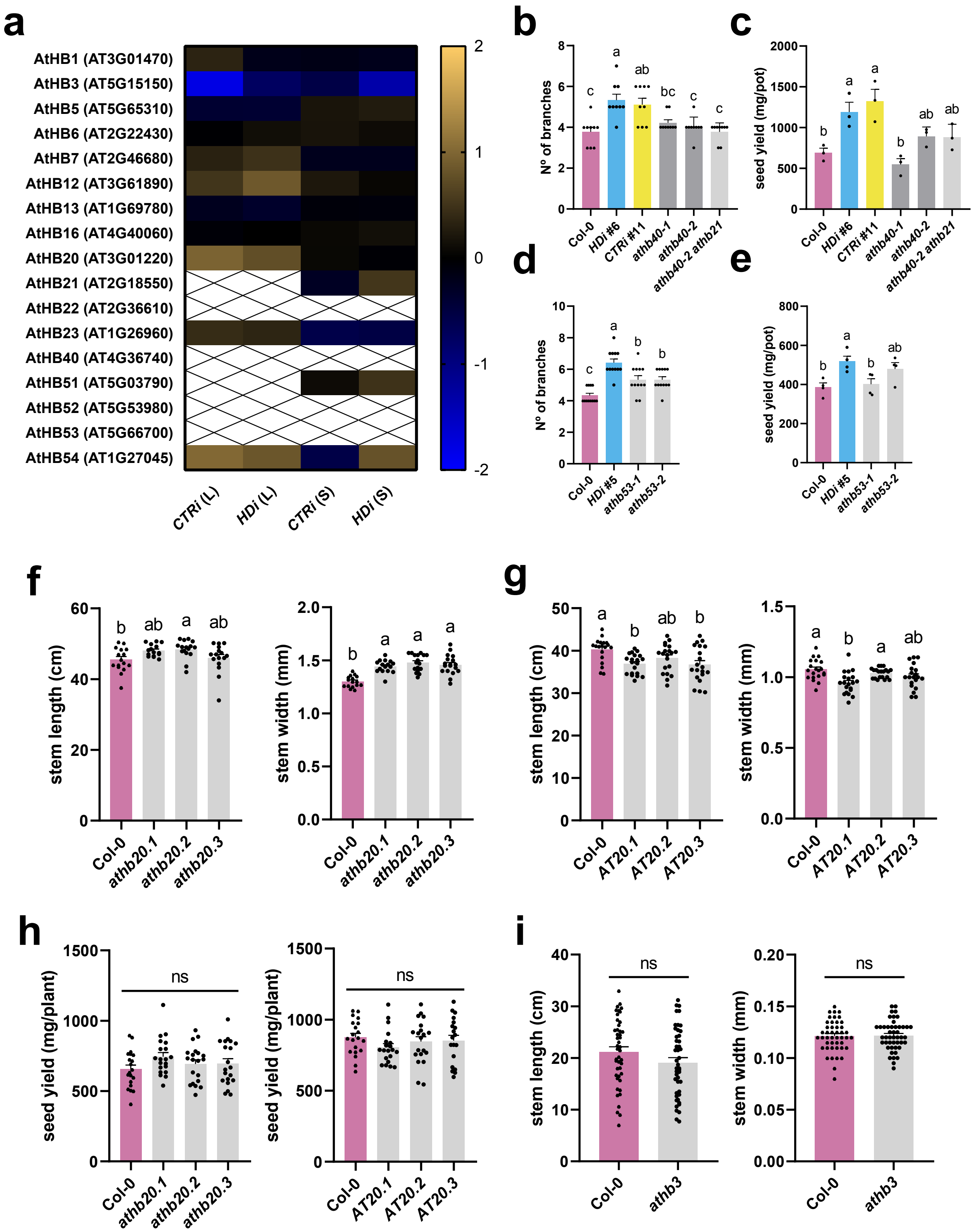


**Figure S4. *AtHB20* and *AtHB53* loss-of-function mutants recapitulate distinct traits of plants expressing *HaHB4*-derived siRNA**

(a) Heatmap of expression changes in all HD-Zip I genes, determined by RNA-seq analysis of leaves (L) and stems (S) of 30-day-old *HDi* or *CTRi* plants. The yellow-to-blue color scale indicates $\log_2$(fold change) relative to the WT level for each given gene. (b) Number of cauline branches of 40-day-old Col-0, *HDi*, *CTRi*, *athb40-1*, *athb40-2*, and *athb40-2 athb21* plants. (c) Seed yield evaluated at the end of the life cycle of the plants described in (b). (d) Number of cauline branches of 40-day-old Col-0, *HDi*, *athb53-1*, and *athb53-2* plants. (e) Seed yield evaluated at the end of the life cycle of the plants described in (d). (f) Stem length (left) and width (right) in Col-0 and three independent athb20 mutant plants (*athb20-1*, *athb20-2*, and *athb20-3*) grown for 36 days. (g) Stem length (left) and width (right) in Col-0 and three independent *AtHB20*-overexpressing lines (*AT20.1*, *AT20.2*, and *AT20.3*) grown for 36 days. (h) Seed yield evaluated at the end of the life cycle of the plants described in (f) and (g). (i) Stem length (left) and width (right) in 36-day-old Col-0 and *athb3* plants. All plants were grown under long-day conditions at 22 °C, with 3 or 4 pots per genotype and three plants per pot. For (c), (e), and (h), seed yield was expressed as mg/pot. All data are means (±SEM) of $n$=3-45 independent biological replicates. Different letters indicate significant differences among means as determined using one-way ANOVA followed by Tukey's *post-hoc* test ($P$<0.05). ns, not significant.

**Table S6.** Set of primers used in this study for cloning.

| **Name** | **Sequence 5' – 3'** | **Restriction site** | **Construct** |
|---|---|---|---|
| RNAi HDS F | GGG CTCGAG ATGTCTCTTCAACAAGTAAC | *Xho*I | pHannibal |
| RNAi HDS R | CCC GAATTC TTGCTCAATCTGCCTCGAC | *Eco*RI | pHannibal |
| RNAi HDAS F | GGG GGATCC ATGTCTCTTCAACAAGTA | *Bam*HI | pHannibal |
| RNAi HDAS R | GGG AAGCTT TTGCTCAATCTGCCTCGAC | *Hind*III | pHannibal |
| RNAi CTRS F | GGG CTCGAG GAGTATAACGCGCTAAAGC | *Xho*I | pHannibal |
| RNAi CTRS R | GCC GAATTC TTAGAACTCCCACCACTTTTG | *Eco*RI | pHannibal |
| RNAi CTRAS F | GGG GGATCC GAGTATAACGCGCTAAAGC | *Bam*HI | pHannibal |
| RNAi CTRAS R | GGG AAGCTT TTAGAACTCCCACCACTTTTG | *Hind*III | pHannibal |

**Table S7.** Set of primers used in this study for genotyping

| Locus | Name | Sequence 5' – 3' |
|---|---|---|
| AT3G49500 | *rdr6 WT F* | CTCTTTTGAGATCATGTTTCTAGT |
| AT3G49500 | *rdr6-12 mut F* | ATCTCTTTTGAGATCATGTTTATT |
| AT3G49500 | *rdr6-12 R* | GAACCATGCAGAGCGGTCTCTCAG |
| | IntronF | TTCTAGCTGGTTTGATGAATTAAATA |
| | IntronR | CAAGCAGATTGGAATTTCTAACAA |

**Table S8.** Set of primers used in this study for RT-qPCR experiments.

| Name | Sequence 5' – 3' |
|---|---|
| HB4qF | GGGCTTCATCCTCGTCAAGTGGC |
| HB4qR | ACGCAAGCGTCTCGTAGTTATG |
| ACTIN2/8 F | GGTAACATTGTGCTCAGTGGTGG |
| ACTIN2/8 R | AACGACCTTAATCTTCATGCTGC |
